# Towards individualized deep brain stimulation: A stereoelectroencephalography-based workflow for neurostimulation target identification

**DOI:** 10.1101/2025.04.22.649607

**Authors:** Jeremy Saal, Kelly Kadlec, Anusha B. Allawala, Lucille Johnston, Ryan B. Leriche, Ritwik Vatsyayan, Yiyuan Han, Audrey Kist, Tommaso Di Ianni, Heather E. Dawes, Edward F. Chang, A Moses Lee, Andrew D. Krystal, Khaled Moussawi, Prasad Shirvalkar, Kristin K. Sellers

**Affiliations:** UCSF Weill Institute for Neurosciences, University of California, San Francisco, San Francisco, CA, USA; Department of Neurological Surgery, University of California, San Francisco, San Francisco, CA, USA; Department of Neurology, University of California, San Francisco, San Francisco, CA, USA; Department of Psychiatry and Behavioral Sciences, University of California, San Francisco, San Francisco, CA, USA; Department of Radiology and Biomedical Imaging University of California, San Francisco, San Francisco, CA, USA; UCSF Department of Anesthesiology and Perioperative Care, Division of Pain Medicine, University of California, San Francisco, San Francisco, CA, USA

**Keywords:** Stereoelectroencephalography, deep brain stimulation, neurostimulation, personalized targets, clinical workflow

## Abstract

**Objectives:** Deep brain stimulation (DBS) is increasingly being used to treat a variety of neuropsychiatric conditions, many of which exhibit idiosyncratic symptom presentations and neural correlates across individuals. Thus, we have utilized inpatient stereoelectroencephalography (sEEG) to identify personalized therapeutic stimulation sites for chronic implantation of DBS. Informed by our experience, we have developed a statistics-driven framework for stimulation testing to identify therapeutic targets.

**Materials and Methods:** Fourteen participants (major depressive disorder = 6, chronic pain = 6, obsessive-compulsive disorder = 2) underwent inpatient testing using sEEG and symptom monitoring to identify personalized stimulation targets for subsequent DBS implantation. We present a structured approach to this sEEG testing, integrating a Stimulation Testing Decision Tree with power analysis and effect size considerations to inform adequately powered results to detect therapeutic stimulation sites with statistical rigor.

**Results:** Effect sizes (Hedges’ g) of stimulation-induced symptom score changes ranged from -1.5 to +2.39. The standard deviation of sham trial responses was a strong predictor of stimulation response variability, as confirmed by a leave-one-out cross-validated linear regression (R^2^ = 0.67, permutation p<0.001). Thus, early sham trial data could be used to estimate the variability of stimulation responses for power analysis calculations. We show that approximately 10 sham trials were needed to robustly estimate sham variability. Power analysis (using a paired-t test) showed that for effect sizes ≥ 1.1, roughly 10 trials should be used per stimulation site for sufficiently powered results.

**Conclusions:** The presented workflow is adaptable to multiple indications and is specifically designed to overcome key challenges experienced during stimulation site testing. Through incorporating sham trials, effect size calculations, and tolerability testing, the described approach can be used to identify personalized and clinically efficacious stimulation sites.

## Introduction

Deep brain stimulation (DBS) is emerging as a promising therapeutic modality for a wide range of indications, particularly neuropsychiatric disorders that involve underlying brain network dysfunctions. Unique perturbations in network function across individuals have led to interest in testing stimulation across brain regions to identify personalized therapeutic sites. To achieve this, our center at the University of California, San Francisco and others utilize inpatient stereoelectroencephalography (sEEG) to explore stimulation across the disease network, aiming to identify personalized therapeutic targets for chronic implantation with DBS (1–8).

Throughout, we use the overarching term “DBS” to refer to both DBS devices and responsive neurostimulation (RNS) using the NeuroPace RNS System. sEEG inpatient monitoring is the established standard of care for complex cases of refractory epilepsy (9,10). We leverage the existing surgical procedures and specialty inpatient units to care for patients with temporarily implanted sEEG leads. We have conducted sEEG monitoring stays lasting approximately 10 days with 14 participants across three clinical indications (major depressive disorder [MDD], chronic pain [CP], and obsessive-compulsive disorder [OCD]). Informed by our experience, we have developed a statistics-driven framework for stimulation testing to identify therapeutic targets.

DBS for complex neuropsychiatric disorders has had variable clinical efficacy across individuals and indications. DBS for MDD has primarily targeted the ventral capsule/ventral striatum (VC/VS) (11) and the subcallosal cingulate (SCC) (12). While smaller open-label studies and case-reports across anatomical targets have shown initial success (13), results from randomized sham-controlled clinical trials have shown considerable heterogeneity (14).

Investigations into DBS for chronic pain date back to the 1950s, revealing potential targets including the anterior cingulate cortex, ventral posterolateral and ventral posteromedial nuclei of the thalamus, and the periventricular grey (PVG) (15,16). Clinical outcomes of DBS for CP are promising, with some studies reporting that over half of patients experience more than 50% pain reduction within 12 months of implantation (17). In the context of OCD, the FDA granted a Humanitarian Device Exemption in 2009 for DBS targeting the anterior limb of the internal capsule. Since then, stimulation has also been targeted to the VC/VS, bed nucleus of the stria terminalis, and subthalamic nucleus (18,19). Yet a third of patients show no response (20), underscoring the need for individualized targeting. Our staged sEEG-DBS approach aims to improve clinical efficacy by personalizing DBS targets.

Current DBS devices support one to four leads, limiting anatomical coverage. Though revision surgeries to reposition leads are possible, they are costly and increase surgical risk. Hence, optimal implantation site (location of stimulation electrode contacts) selection is crucial, and using sEEG allows for investigating the same number of targets that would require the equivalent of three to five revision DBS surgeries to evaluate. Towards this objective, our testing framework is designed with three goals in mind: (1) standardize stimulation procedures for rigorous results; (2) minimize risk and burden caused by the failure of DBS surgery and/or revision surgery to find therapeutically beneficial stimulation sites; and (3) identify at least one robust therapeutic target.

To address these goals, we present here a Stimulation Testing Decision Tree designed to facilitate the identification of personalized stimulation sites. Importantly, the Decision Tree incorporates power analysis and effect size considerations based on data from participants who underwent stimulation testing with sEEG in 3 indications (MDD, CP, OCD), to inform the design of stimulation testing which will be adequately powered to detect therapeutic stimulation sites with statistical rigor. We propose this testing framework to avoid pitfalls and obstacles that we have identified during prior sEEG testing periods, including:

1. Trying to (over)optimize one site of stimulation: Without a principled testing approach, upon observing therapeutic benefit with stimulation, it can be tempting to adjust stimulation parameters to try and ‘optimize’ stimulation at that site. However, this is often done at the expense of obtaining sufficient trials with stable parameters to statistically power a determination of benefit, testing other sites, or conducting sufficient sham-trial testing.
2. Over-reliance or under-reliance on quantitative metrics to assess therapeutic benefit: Even with training on self-report measures, participants adopt idiosyncratic behavior in responding to symptom surveys, such as only responding within a narrow range of the scale. Thus, over-reliance on strict quantitative cutoffs may not fully capture beneficial stimulation. Conversely, basing surgical implant decisions entirely on clinician observations may suffer from personal bias and inter-rater reliability concerns.
3. Weighing stimulation benefits and side effects: Even if stimulation is highly beneficial for symptom relief, if it is accompanied by concerning side effects, other sites should be pursued for benefit. Stimulation-induced side effects may prevent participant blinding during randomized controlled trials. These side effects span a range of experiences, including sweaty hands, buzzing or tingling, bladder spasms, dissociation, vertigo, induced smiling, and others.

## Materials and Methods

### Study Participants and Clinical Environment

As part of respective clinical trials, 14 participants were implanted with sEEG for approximately 10 days of inpatient monitoring to identify brain regions in which acute stimulation led to reduction in primary indication symptoms (MDD: n = 6, CP: n = 6, OCD: n = 2). PMT (PMT Corporation, Chanhassen, MN), AdTech (AdTech Medical, Oak Creek, WI), or DIXI (DIXI Medical, Marchaux Chaudefontaine, France) electrodes were implanted bilaterally in regions implicated in the primary indication. Anatomical localization of electrode contacts was conducted by co-registration of preoperative T1 MRI images and postoperative CT scan and automated parcellation and manual verification of the labeling. Contacts within defined regions of interest were selected for subsequent stimulation testing. Electrode tails were connected to cables that interfaced with a 256-channel Nihon Kohden neurophysiology recording and stimulation system. Charge-balanced, biphasic, bipolar stimulation was delivered between one anode and one cathode. The duration of stimulation used for testing ranged across indications based on expected and observed modulation of symptoms (10s to 20 minutes of continuous stimulation).

Participants provided frequent symptom reports via visual analog scale (VAS) ratings on a tablet computer. VAS sliders were horizontal and ranged from 0 (no presence of that symptom) to 100 (highest presence of that symptom). VAS symptoms were indication-specific (MDD: depression; CP: pain intensity; OCD: obsessions, compulsions, and OCD-related distress [scores were summed to create an OCD composite score]). Scores were collected before stimulation trials (baseline), during stimulation or sham trials, and during washout. These data were collected and managed using REDCap (Research Electronic Data Capture) hosted at University of California, San Francisco (21,22). REDCap is a secure, web-based software platform designed to support data capture for research studies providing 1) an intuitive interface for validated data capture; 2) audit trails for tracking data manipulation and export procedures; 3) automated export procedures for seamless data downloads to common statistical packages; and 4) procedures for data integration and interoperability with external sources. Change in VAS score (score during stimulation/sham minus score during the immediately preceding baseline or washout period, termed ‘change score’) was used to quantitatively assess for symptom changes induced by stimulation. We focus our analysis on VAS survey responses because of their utility for momentary symptom tracking. All study activities were approved by the Committee on Human Research at the University of California, San Francisco and covered under FDA Investigational Device Exemptions for each indication (MDD: NCT04004169, CP: NCT04144972, OCD: NCT06347978).

### Statistical Analysis

For the retrospective analysis of effect size on data which lacked a paired trial structure, we used Hedges’ g for independent groups.

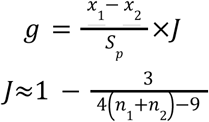

where,

● 𝑥_1̱_ and 𝑥_2̱_ are the means of the two groups
● 𝑛_1_ and 𝑛_2_ are the sample sizes of the two groups
● 𝑆_*p*_ is the pooled standard deviation of the two groups

A key advantage of the proposed workflow is the paired, block-based design (Supplemental Table 1) which will enable paired analyses in future data sets, such as paired t-tests and Cohen’s d_z_ for effect size.

Cohen’s d_z_ is calculated using the formula

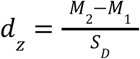

where 𝑆_*D*_ is the standard deviation of the differences, calculated as:

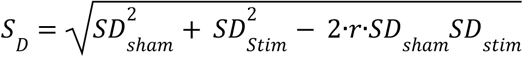

where 𝑆𝐷*_sham_* and 𝑆𝐷*_sham_* are the standard deviations of the sham and stimulation data, respectively, and 𝑟 is the correlation between sham and stimulation data.

One component of the proposed workflow is determining the required number of trials per site through a prospective power analysis. Prospectively calculating this effect size for a power analysis requires estimates for two unknown parameters: the correlation (r) between paired sham and verum stimulation trial change scores and their response variabilities (SDs). We recommend assuming a conservative value for the correlation (e.g. 0.2) to avoid overestimating the effect size. Estimating the SDs is more difficult because the identity of the best stimulation site is unknown. We hypothesized that the SD from sham trial difference scores could serve as a reliable predictor for SD of change scores at the best stimulation site.

To quantify whether change score (the symptom score during a trial minus the score from the immediately preceding baseline) variability in sham trials is correlated with that of verum stimulation at sites with large effect sizes, we calculated the Pearson correlation. Further, to test whether variability from sham trials could be used to estimate SDs for responses to the best verum stimulation sites, we used a leave-one-out (LOO) cross-validated linear regression. For each of the 14 participants, a linear regression model was trained on the remaining 13 participants and used to predict the held-out participant’s best-site stimulation change score SD from their sham change score SD. The statistical significance of the resulting R^2^ value was assessed using a permutation test in which stimulation response change score SDs were randomly shuffled 5,000 times before the LOO regression was re-calculated; the true R^2^ value was then compared to the resulting null distribution.

### Number of Trials Calculation

To estimate the required sample size at different clinically relevant effect sizes (0.8 to 1.7) to achieve sufficiently powered results (power = 0.8), we used the open-source computing tool G*Power (23). Due to the invasive nature of the study, we are focusing on large effect sizes. Furthermore, this range was chosen to reflect effect sizes observed (Fig 2A). Given the paired design proposed for future studies, we conducted this power analysis using a paired one-tailed t-test with a power of 0.8 and alpha = 0.05 (one-tailed because the desired effect is in one direction only). Bonferroni multiple comparisons correction was applied to this calculation to determine the required sample size when multiple regions were tested (e.g. alpha = 0.05 for 1 region; alpha = 0.005 for 10 regions). A similar procedure was followed using unpaired t-tests and paired Wilcoxon signed rank to assess sample sizes for unpaired and nonparametric distributions, respectively.

**Figure 1.**
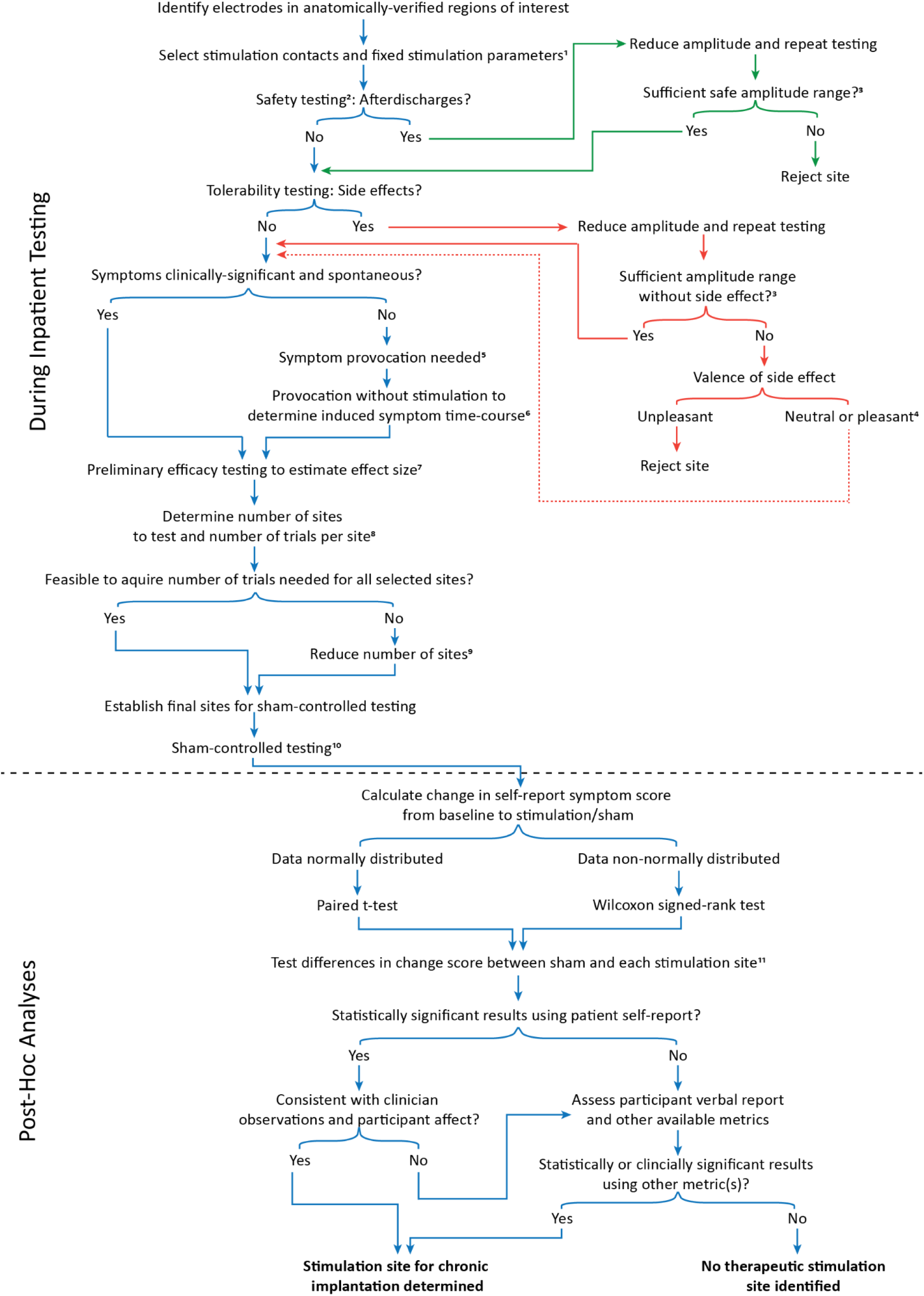
The Stimulation Testing Decision Tree provides a systematic and statistical approach for identifying therapeutic stimulation sites. 1: The parameter space is too large to exhaustively explore during inpatient testing. Select contact pairs for bipolar stimulation (anode and cathode) within each region of interest (ROI) based on co-registration of MRI and CT images. If multiple electrode contacts are within an ROI, different contact pairs can be tested for safety and tolerability. To manage the large number of possible stimulation parameters, we recommend initially holding frequency and pulse-width constant. This pragmatic trade-off increases statistical power for determining a stimulation location, though a new parameter set at the same anatomical target can simply be treated as a new ’site’ for comparison. 2: Initial safety testing recommendation: 1s and then 3s at 1, 3, and 6mA using selected fixed stimulation parameters, per electrode contact pair (these parameters may need to be adjusted if using electrodes with small contact sizes or non-standard pitch). 3: Sufficient amplitude range may differ between cortical and subcortical targets. For both safety and side effects, we recommend rejecting sites if stimulation below 1mA for cortical and 0.8mA for subcortical sites is necessary. These amplitude ranges may need to be adjusted if using electrodes with small contact sizes or non-standard pitch. 4: The presence of side effects with stimulation may compromise participant blinding and subsequent double-blind randomized controlled trials. 5: Symptom provocation may include discussion of triggering life events (MDD), exposure to cues (OCD), or physical movement (CP). The specific activity should be tailored to a given participant’s symptom profile and presentation. 6: Symptoms elicited by provocation may dissipate; the time-course of evoked symptoms should be measured and then monitored to inform when additional provocation instances are required. 7: A subset of stimulation sites and parameters should be selected from those which survived safety and tolerability testing for preliminary efficacy testing. Estimated effect size is based on both the sham trial standard deviation as well as a change score value that indicates a clinical level of improvement. 8: The number of sites which can be tested should be determined by using Fig 2, which provides sample size based on estimated effect size and number of sites. 9: If the number of trials required for the sites that remain after safety and tolerability testing is not feasible for testing, the number of sites to be tested must be reduced. See main text for examples of methods to reduce the number of test sites. 10: Testing should be conducted using blocks within the same session consisting of at least one sham trial and up to one trial at each stimulation site. The sham trial is considered to be paired with each of the stimulation sites tested in that block. If multiple trials of sham are tested within one block, the scores should be averaged. See Supplemental Table 1. 11: Correct for multiple comparisons with, for example, Bonferroni’s method.

**Figure 2.**
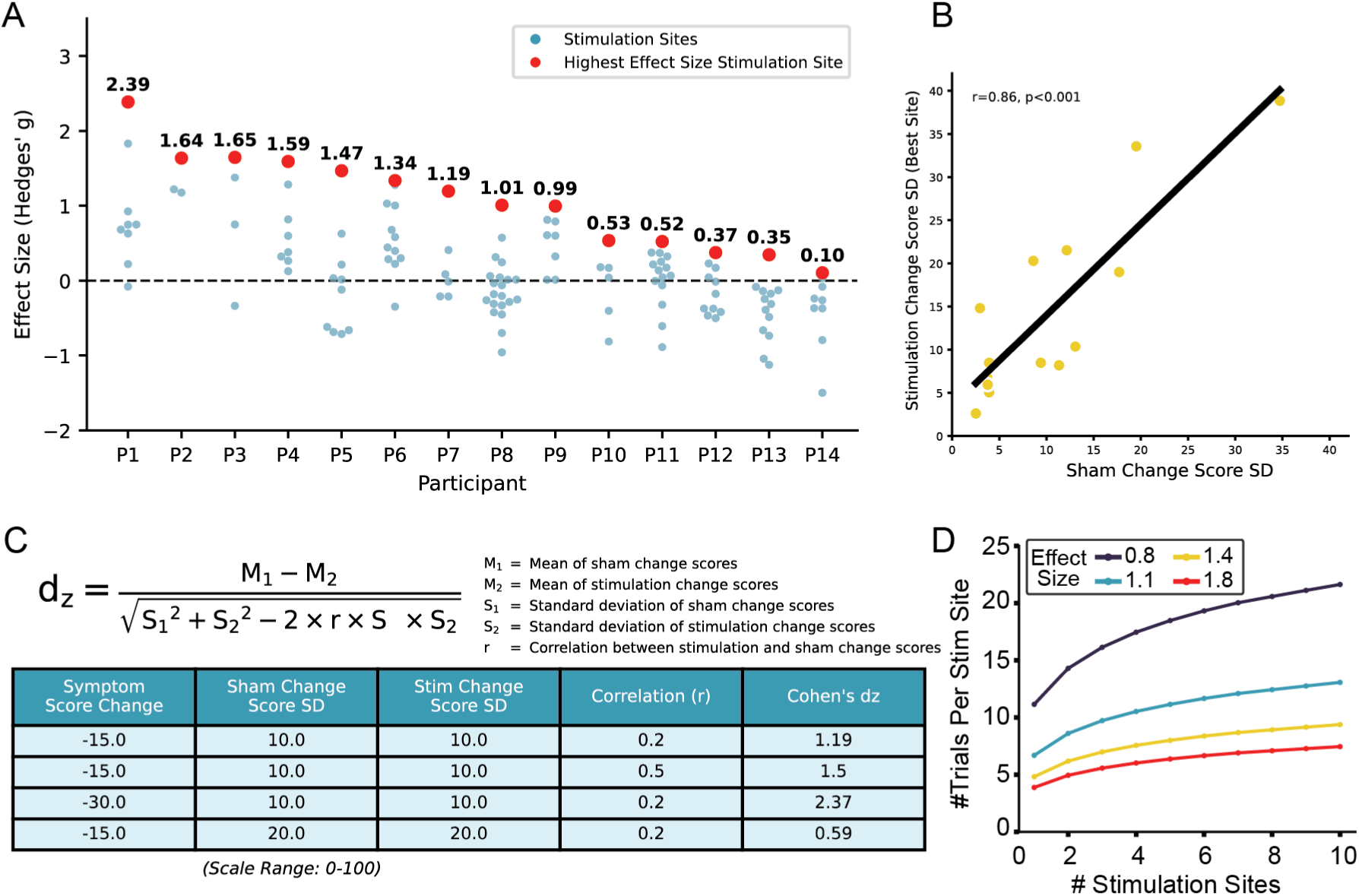
Effect Size Across Participants and Sample Size Estimation for Statistical Power. **A:** Hedges’ g (verum stimulation vs. sham) for all stimulation sites tested in P1-P14. Red points indicate each participant’s largest positive effect size, and blue points indicate effect sizes for all other stimulation sites tested. The dashed line at g = 0 represents no difference from sham. Hedges’ g > 0 indicates improved symptoms relative to sham, whereas g < 0 indicates symptom worsening. **B:** Strong positive correlation between each participant’s sham change score SD and the change score SD at their best stimulation site (r = 0.86, p < 0.001). Leave-one-out linear regression further supported that sham change score SD is a strong predictor of best-site verum stimulation change score SD (R^2^ = 0.67, permutation p<0.001). **C:** Cohen’s d_z_ formula and corresponding variables. The table provides three example calculations. **D:** Number of trials required for statistical significance as a function of the number of brain regions, based on a paired t-test. Each line corresponds to a different effect size. Larger effect sizes require fewer trials to reach significance, while smaller effect sizes require more trials.

### Stability of sham standard deviation across trials

To inform how many preliminary trials are needed to estimate change score response variability, we sought to determine the minimum number of sham trials required for a stable estimation of the full sample change score SD. To this end, we performed a bootstrap analysis on the sham change score SDs after removing outliers (1.5 times interquartile range), starting with two trials and increasing up to the total number of sham trials (10,000 bootstraps, trials sampled with replacement). We then computed the 95% confidence interval (CI) of the distribution of 10,000 SD values. Finally, we examined the half-width of this CI as a representation of the precision of our SD values for estimates using each number of trials.

Lastly, to test if sham change score SD remained stable across the sEEG testing period, we compared the change score SDs of the first and second halves of temporally ordered trials using a paired t-test.

## Results

### Stimulation Testing Decision Tree for Identifying a Therapeutic Stimulation Site

Using data collected from sEEG monitoring phases with 14 participants, we formalized a Stimulation Testing Decision Tree (Fig 1). The Decision Tree can be followed for testing an individual participant and is designed to optimize decision-making based on positive evidence while taking into consideration key points of identified risk. The key parameter we aim to ‘map’ during inpatient testing is the therapeutic stimulation site. Stimulation intensity is adjusted as needed during testing. To focus on maintaining statistical power for determining an efficacious stimulation location, we recommend initially holding frequency and pulse-width constant. Any adjustments to these parameters at the same anatomical target can then be treated as a distinct ’site’ for statistical analysis. The different color-coded sections of the decision tree reflect key considerations to reduce biases in decision making about where to implant chronic DBS for therapeutic benefit.

1. **Evidence for therapeutic efficacy** (Fig 1, blue):

a. *Safety testing*: All stimulation sites should first undergo safety testing. We recommend 1s and then 3s at 1, 3, and 6mA (if using electrodes with 2mm contacts and 0.8mm diameter). If afterdischarges are observed, see ‘Sufficient safe dynamic range’ section below.
b. *Tolerability testing*: Ideally, stimulation does not elicit side effects. If any somatic, sensory, or behavioral side effects are reported or observed with stimulation testing, see ‘Tolerability and side effects of stimulation’ section below.
c. *Spontaneous vs. provoked symptoms*: If symptoms are not spontaneously present, symptom provocation may be required before stimulation testing can continue. Symptom provocation should be thoroughly discussed with participants prior to initiation, including allowable or non-allowable topics of discussion or tasks. While it remains poorly understood if there is a shared neurobiological substrate for elicited vs. spontaneous symptoms, our experience is that stimulation shown to be acutely beneficial in the inpatient context of symptom provocation for MDD and OCD is also beneficial during chronic DBS in the outpatient setting (8).
d. *Symptom baseline:* To effectively test for improvement, stimulation trials should only proceed when a participant’s spontaneous symptoms are present above a predetermined, clinically meaningful threshold. If symptoms are below this threshold, testing should be paused until they naturally increase or until provocation is used as described in the previous step.
e. *Variability and Effect Size Estimation*: To facilitate designing stimulation testing blocks with statistical power, the expected effect size should first be estimated. Preliminary stimulation blocks can be used to determine the expected variability of symptom responses to stimulation (see Supplementary Table 1 for suggested trial design). Based on our data, variability from sham trials is a strong predictor of the variability of the best stimulation site (R^2^ = 0.67, permutation p<0.001). Therefore, an initial set of stimulation blocks should include a higher density of sham trials compared to subsequent efficacy testing. This provides the robust data needed to estimate the effect size before finalizing the number of sites for powered testing.
f. *Determine the number of sites and number of trials per site for testing*: The estimated effect size from the prior step can then be used to determine the number of trials required per stimulation site being tested.
g. *Select stimulation sites*: If there are too many sites remaining after safety and tolerability testing to feasibly acquire the necessary number of trials per site, the number of sites to be tested should be reduced. This can be done using methods appropriate for the particular indication or participant, for example examining tractography, using evoked potentials to evaluate connectivity, performing further stimulation testing to identify sites with strong/rapid behavioral responses, or using prior literature.
h. *Sham-controlled testing*: Stimulation parameters should follow what was determined to be safe and tolerable during safety testing.
i. *Calculate symptom change scores*: A symptom score will be taken immediately preceding and following each stimulation or sham trial to assess changes in symptoms.
j. *Statistical tests*: Either a paired t-test or a paired signed rank test can be used to determine the therapeutic effect of a stimulation site compared with sham.
k. *Results interpretation*: If the statistical tests demonstrate significance for a site compared to sham and this is consistent with clinical observation and participant verbal report, the site can be considered a potential therapeutic target for chronic implantation. If multiple sites are identified as being therapeutic, the most effective site can be determined by ranking therapeutic target change scores. If the statistical test is not significant other metrics (e.g. participant verbal reports, clinician scales) should be investigated for evidence of therapeutic benefit (based on our experience, discrepancies between statistical tests and clinician observations occur in about 15% of cases). Responses should be assessed per individual to determine meaningful clinical benefit. If no clinical benefit is captured by these metrics, no therapeutic site has been identified.
2. **Sufficient safe dynamic range** (Fig 1, green): The presence of afterdischarges with stimulation should warrant extra vigilance at that site. However, safe stimulation may be possible at lower amplitudes (e.g. afterdischarges seen at 6mA do not preclude safe stimulation at 2mA). The threshold for afterdischarges is dynamic, so further stimulation with lower amplitude at a site that exhibited afterdischarges should be undertaken with caution.
3. **Tolerability and side effects of stimulation** (Fig 1, orange): Stimulation-induced side effects can generally be categorized as unpleasant or neutral/pleasant. For example, sensory thalamus stimulation in patients with chronic pain has been shown to induce pleasant paresthesias (24) and PVG stimulation has coincided with a warm sensation (25,26). Unpleasant side effects must be avoided, and the site only remains viable if the side effect is not present with lowered stimulation amplitude. Unwanted behavioral side effects should also be avoided, for example stimulation-evoked anger or mania. The threshold for side effects is dynamic, so sufficient amplitude range without side effects must be present during acute testing. If participant blinding is necessary, as is the case for many clinical trials, sites should only be advanced if amplitude can be lowered to the point of no perceptible side effects, even if side effects are neutral or positive. If blinding is not needed, clinicians/researchers should carefully evaluate together with participants if the presence of neutral or positive side effects would be distracting or maladaptive in daily life. Another consideration in evaluating tolerability is that some side effects, in particular those that are transient and correspond to stimulation onset/offset, may be present during testing but can be avoided in the outpatient setting.

### Empirical determination of sample size

Across all participants and all stimulation sites, Hedges’ g values ranged from −1.5 to +2.39 (Fig 2A). We were interested in the greatest positive effect size stimulation site within each participant, which indicates an improvement in symptoms due to stimulation (range of 0.1 to 2.39 across the 14 participants). Each participant’s sham change score SD versus their best-site change score SD revealed a strong positive relationship (r = 0.86, p < 0.001), as illustrated in Figure 2B. Participants with higher variability in sham trials (e.g., those whose sham SD exceeded 20) tended to have equally large best-site variability (e.g., 30+), suggesting trait-level characteristics in the variability of score reporting. To more robustly test the predictive power of this relationship, we performed a leave-one-out cross-validated regression. This analysis confirmed that sham change score SD was a strong out-of-sample predictor of best-site stimulation change score SD (R^2^ = 0.67, p< 0.001). Thus, we can assume a similar SD of changes scores for both the sham and verum stimulation trials. Leveraging this fact, change scores from sham from preliminary stimulation testing blocks can be used to estimate the SD of the best stimulation site (detailed in block trial design below). This estimate can then be used to estimate the expected Cohen’s d_z_ (Figure 2C).

Using the statistical power analysis tool G*power (22), we computed the number of trials required for different effect sizes and numbers of stimulation sites. For Cohen’s d_z_ ≥ 1.1, around 10 samples per site are needed for statistically powered results (Fig 2D). The same analysis performed for the case of an unpaired t-test shows a substantial increase in the number of samples required per site, emphasizing the importance of a paired design with matched trials for sham and stimulation site (Supplemental Fig 1A). This increase in required samples per site does not occur when using a paired nonparametric test for non-normally distributed data (Supplemental Fig 1B).

We propose a blocked trial design in which each block has at least one sham trial plus any number of different verum stimulation sites within a given testing session (Supplemental Table 1). Before sham-controlled efficacy testing, preliminary testing blocks with a higher concentration of sham trials can provide an estimate for sham change score SD and thus effect size. To determine the number of preliminary sham trials needed to robustly estimate the change score SD, we performed a bootstrap analysis on data from three participants with a sufficient number of sham trials. Of these, two participants each had a single statistical outlier trial which were removed from this analysis. For participants with low and moderate variability, the change score SD estimate improved rapidly and then began to level-off after around 10 trials (Supplemental Figure 1C), whereas the participant with high variability scores did not reach a plateau.

Of important note is that even small changes in SD can have a large impact on effect size calculations. Bootstrap analysis can be used to generate an upper and lower bound for SD estimates to inform a more conservative estimate for effect size calculation. As a final confirmation of the stability of sham change score SD across days, a paired-samples t-test revealed no significant difference in sham response variability between the first and second half of trials (t(13) = 0.756, p = 0.463), supporting the stability of sham trial variability for estimating stimulation response SD (Supplemental Fig 1D). This said, the results of our bootstrap analysis illustrate a known characteristic of SD which is that acquiring more samples will allow for more precise estimation. Thus, it is important that as sham-controlled testing progresses, the SD is continuously evaluated to ensure adjustments to expected effect size are made if necessary.

## Discussion

We propose an empirically derived and generalizable sEEG-based workflow to identify therapeutic sites for chronic implantation of DBS electrodes to treat a variety of neuropsychiatric conditions. Using lessons learned from 14 participants undergoing a sEEG trial period, this workflow involves clear decision-making rationale for both inpatient testing and post-hoc analyses. Our sEEG-based decision tree and workflow offer a prospective method to systematically reduce bias while enhancing both personalization and standardization in chronic DBS implantation site selection. By leveraging trait-level variability in symptom score reporting, we can estimate effect sizes from sham trials during preliminary stimulation testing and determine the number of trials needed per stimulation site to provide statistically powered results.

One foundational assumption of our sEEG-based workflow is that acute, stimulation-induced symptom improvements will predict long-term clinical outcomes from chronic deep brain stimulation. Prior work has demonstrated that acute stimulation during a stereo-EEG period does lead to sustained benefit in treatment-resistant depression, chronic pain, and obsessive-compulsive disorder (3,6,7,8). However, it is important to note that our methodology is used to seek targets where acute stimulation is a sufficient condition for therapeutic benefit. The proposed workflow may miss effective sites that require longer periods of chronic stimulation (27). Therefore, our workflow should be considered a tool for identifying rapid-response brain stimulation targets, rather than an exhaustive survey of all potential therapeutic sites.

Despite its advantages, this workflow has several limitations. For instance, microlesion effects caused by the implant and/or explant surgeries can alter symptom intensity and may contribute to changes observed during inpatient mapping. It is important to monitor symptoms after the sEEG leads are removed; a persistent improvement in symptoms may indicate a microlesion effect, which should be taken into account when considering permanent DBS implantation. While sham trials and washout periods help mitigate carryover concerns, available time may still be insufficient for complete symptom reversion. Untested parameters at a “ruled-out” site could theoretically outperform the chosen implantation site. The proposed framework does not guarantee the identification of a therapeutic target. Rather, it provides testing guidelines which, based on real participant data, are likely to provide sufficient statistical power to identify a therapeutic target if it exists within the regions tested. In future studies, data-driven methods based on Bayesian optimization or machine learning models could be integrated with our framework to effectively optimize over a larger parameter space (28–30). Additionally, this framework does not specifically facilitate the assessment of state-dependent effects of stimulation, as testing is aggregated across multiple days and states.

While these guidelines are designed to be generalized as they are based on multiple indications, it is important to note that this is limited by the small number of participants and will have to be evaluated in larger studies. Long-term outcomes from the workflow are also not yet known but will be critical to evaluate when blinded testing with chronically implanted devices is concluded. Another limitation of this workflow is the current reliance on VAS scales as the main measure of therapeutic effect. Although VAS scales have been shown to correspond with validated clinical measures (31,32), there is still a need for improved measurements of acute fluctuations in symptoms.

Use of this workflow will provide statistically rigorous results which can improve future sEEG targeting and testing. To this end, we are currently working to understand the mechanisms engaged by therapeutic vs non-therapeutic stimulation by combining models of the volume of activated tissue with tractography to estimate the white matter tracts implicated in stimulation. This will be a critical part of further personalizing target selection. It is worth noting that although this workflow provides needed guidelines for standardizing testing in participants undergoing sEEG ahead of chronic DBS implant, this type of testing may not be scalable if DBS becomes a more widely adopted treatment for psychiatric conditions. For this reason, it will also be critical to extend the neurophysiological insight gained from this method to nonsurgical methods in the future.

## Conclusion

To summarize, we have proposed a data-driven workflow that leverages sEEG-based stimulation to address key challenges in personalizing DBS. The proposed approach allows the identification of therapeutic sites by integrating rigorous trial design, effect size analysis, and sham comparisons. While this approach should be further validated, our framework provides a pragmatic method for optimizing DBS targets and parameters across diverse neuropsychiatric conditions, thus laying the groundwork for more precise and effective neuromodulation therapies.

## Authorship Statements

JS, KK, ABA, LJ, RBL, RV, YH, AK, TDI, HED, EFC, AML, ADK, KM, PS, and KKS designed and conducted the studies, including patient recruitment, data collection, data analysis, and interpretation of the data. JS, KK, RV, YH, KKS prepared the manuscript draft with intellectual input from ABA, LJ, AK, TDI, HED, AML, ADK, KM, and PS. All authors approved the final manuscript

## References

1. Starkweather CK, Sugrue LP, Cajigas I, Speidel B, Krystal AD, Scangos K, Chang EF (2024): Stereoelectroencephalography Electrode Implantation for Inpatient Workup of Treatment-Resistant Depression. Neurosurgery 95: 941–948.

2. Shirvalkar P, Sellers KK, Schmitgen A, Prosky J, Joseph I, Starr PA, Chang EF (2020): A Deep Brain Stimulation Trial Period for Treating Chronic Pain. J Clin Med 9: 3155.

3. Scangos KW, Khambhati AN, Daly PM, Makhoul GS, Sugrue LP, Zamanian H, et al. (2021): Closed-loop neuromodulation in an individual with treatment-resistant depression. Nat Med 27: 1696–1700.

4. Sheth SA, Shofty B, Allawala A, Xiao J, Adkinson JA, Mathura RK, et al. (2023): Stereo-EEG-guided network modulation for psychiatric disorders: Surgical considerations. Brain Stimulat 16: 1792–1798.

5. Allawala A, Bijanki KR, Goodman W, Cohn JF, Viswanathan A, Yoshor D, et al. (2021): A Novel Framework for Network-Targeted Neuropsychiatric Deep Brain Stimulation. Neurosurgery 89: E116–E121.

6. Sheth SA, Bijanki KR, Metzger B, Allawala A, Pirtle V, Adkinson JA, et al. (2022): Deep Brain Stimulation for Depression Informed by Intracranial Recordings. Biol Psychiatry 92: 246–251.

7. Shirvalkar P, Leriche R, Saal J, Cagle J, Prosky J, Joseph I, et al. (2025): Personalized, closed-loop deep brain stimulation for chronic pain. *medRxiv*.

8. Lee AM, Kist A, Alvarez J, Sellers KK, Khambhati AN, Sugrue LP, et al. (2025): Invasive Brain Mapping Identifies Personalized Therapeutic Neuromodulation Targets for Obsessive-Compulsive Disorder. *medRxiv*.

9. Theodore WH, Fisher RS (2004): Brain stimulation for epilepsy. Lancet Neurol 3: 111–118.

10. Delgado-Garcia G, Frauscher B (2022): Future of Neurology & Technology: Stereoelectroencephalography in Presurgical Epilepsy Evaluation. Neurology 98.

11. Malone DA, Dougherty DD, Rezai AR, Carpenter LL, Friehs GM, Eskandar EN, et al. (2009): Deep Brain Stimulation of the Ventral Capsule/Ventral Striatum for Treatment-Resistant Depression. Biol Psychiatry 65: 267–275.

12. Lozano AM, Mayberg HS, Giacobbe P, Hamani C, Craddock RC, Kennedy SH (2008): Subcallosal Cingulate Gyrus Deep Brain Stimulation for Treatment-Resistant Depression. Biol Psychiatry 64: 461–467.

13. Hitti FL, Cristancho MA, Yang AI, O’Reardon JP, Bhati MT, Baltuch GH (2021): Deep Brain Stimulation of the Ventral Capsule/Ventral Striatum for Treatment-Resistant Depression: A Decade of Clinical Follow-Up. J Clin Psychiatry 82.

14. Dougherty DD, Rezai AR, Carpenter LL, Howland RH, Bhati MT, O’Reardon JP, et al. (2015): A Randomized Sham-Controlled Trial of Deep Brain Stimulation of the Ventral Capsule/Ventral Striatum for Chronic Treatment-Resistant Depression. Biol Psychiatry 78: 240–248.

15. Shirvalkar P, Veuthey TL, Dawes HE, Chang EF (2018): Closed-Loop Deep Brain Stimulation for Refractory Chronic Pain. Front Comput Neurosci 12: 18.

16. Boccard SGJ, Pereira EAC, Aziz TZ (2015): Deep brain stimulation for chronic pain. J Clin Neurosci 22: 1537–1543.

17. Farrell SM, Green A, Aziz T (2018): The Current State of Deep Brain Stimulation for Chronic Pain and Its Context in Other Forms of Neuromodulation. Brain Sci 8: 158.

18. De Koning PP, Figee M, Van Den Munckhof P, Schuurman PR, Denys D (2011): Current Status of Deep Brain Stimulation for Obsessive-Compulsive Disorder: A Clinical Review of Different Targets. Curr Psychiatry Rep 13: 274–282.

19. Mosley PE, Windels F, Morris J, Coyne T, Marsh R, Giorni A, et al. (2021): A randomised, double-blind, sham-controlled trial of deep brain stimulation of the bed nucleus of the stria terminalis for treatment-resistant obsessive-compulsive disorder. Transl Psychiatry 11: 190.

20. Gadot R, Najera R, Hirani S, Anand A, Storch E, Goodman WK, et al. (2022): Efficacy of deep brain stimulation for treatment-resistant obsessive-compulsive disorder: systematic review and meta-analysis. J Neurol Neurosurg Psychiatry 93: 1166–1173.

21. Harris PA, Taylor R, Thielke R, Payne J, Gonzalez N, Conde JG (2009): Research electronic data capture (REDCap)—A metadata-driven methodology and workflow process for providing translational research informatics support. J Biomed Inform 42: 377–381.

22. Harris PA, Taylor R, Minor BL, Elliott V, Fernandez M, O’Neal L, et al. (2019): The REDCap consortium: Building an international community of software platform partners. J Biomed Inform 95: 103208.

23. Faul F, Erdfelder E, Lang A-G, Buchner A (2007): G*Power 3: A flexible statistical power analysis program for the social, behavioral, and biomedical sciences. Behav Res Methods 39: 175–191.

24. Pereira EAC, Boccard SG, Linhares P, Chamadoira C, Rosas MJ, Abreu P, et al. (2013): Thalamic deep brain stimulation for neuropathic pain after amputation or brachial plexus avulsion. Neurosurg Focus 35: E7.

25. Young RF, Chambi VI (1987): Pain relief by electrical stimulation of the periaqueductal and periventricular gray matter: Evidence for a non-opioid mechanism. J Neurosurg 66: 364–371.

26. Richardson DE, Akil H (1977): Pain reduction by electrical brain stimulation in man. J Neurosurg 47: 178–183.

27. Himes LM, Mayberg HS, Husain MM, Holtzheimer PE, Lozano AM, Kennedy SH, et al. (2025): Revisiting subcallosal cingulate deep brain stimulation for depression: Long-term safety and effectiveness outcomes from a pooled analysis of 172 implanted patients. Brain Stimulat 18: 1632–1640.

28. Stieve BJ, Richner TJ, Krook-Magnuson C, Netoff TI, Krook-Magnuson E (2023): Optimization of closed-loop electrical stimulation enables robust cerebellar-directed seizure control. Brain 146: 91–108.

29. Minai Y, Soldado-Magraner J, Smith MA, Yu BM (2024): MiSO: Optimizing brain stimulation to create neural population activity states. Advances in Neural Information Processing Systems 37. presented at the NeurIPS 2024.

30. Boutet A, Madhavan R, Elias GJB, Joel SE, Gramer R, Ranjan M, et al. (2021): Predicting optimal deep brain stimulation parameters for Parkinson’s disease using functional MRI and machine learning. Nat Commun 12: 3043.

31. Lingjærde O, Føreland AR (1998): Direct assessment of improvement in winter depression with a visual analogue scale: high reliability and validity. Psychiatry Research 81: 387–392.

32. Amin MR, Siratinayer M, Abadi A, Moradyan T, (2012): Correlation between visual analogue scale and short form of McGill questionnaire in patients with chronic low back pain. Qom Univ. Med. Sci. J 5, 31–34.

