## Supplemental Files for "Towards individualized deep brain stimulation: A stereoelectroencephalography-based workflow for neurostimulation target identification"

Running Title: Workflow for stimulation target discovery

Jeremy Saal MS*[a,b], Kelly Kadlec PhD*[a,c,d], Anusha B. Allawala PhD[a,b], Lucille Johnston BS[a,b], Ryan B. Leriche BS[a,b], Ritwik Vatsyayan PhD[a,b], Yiyuan Han PhD[a,b], Audrey Kist PhD[a,d], Tommaso Di Ianni PhD[a,d,e], Heather E. Dawes PhD[a,b], Edward F. Chang MD[a,b], A Moses Lee MD PhD[a,d], Andrew D. Krystal MD[a,d], Khaled Moussawi MD PhD[a,c,d], Prasad Shirvalkar MD PhD[a,b,c,f], Kristin K. Sellers PhD[a,b]

a. UCSF Weill Institute for Neurosciences, University of California, San Francisco, San Francisco, CA, USA.
b. Department of Neurological Surgery, University of California, San Francisco, San Francisco, CA, USA.
c. Department of Neurology, University of California, San Francisco, San Francisco, CA, USA.
d. Department of Psychiatry and Behavioral Sciences, University of California, San Francisco, San Francisco, CA, USA.
e. Department of Radiology and Biomedical Imaging University of California, San Francisco, San Francisco, CA, USA.
f. UCSF Department of Anesthesiology and Perioperative Care, Division of Pain Medicine, University of California, San Francisco, San Francisco, CA, USA.
* These authors contributed equally

**Contents:**

- **Supplementary Figure 1**
- **Supplementary Table 1**

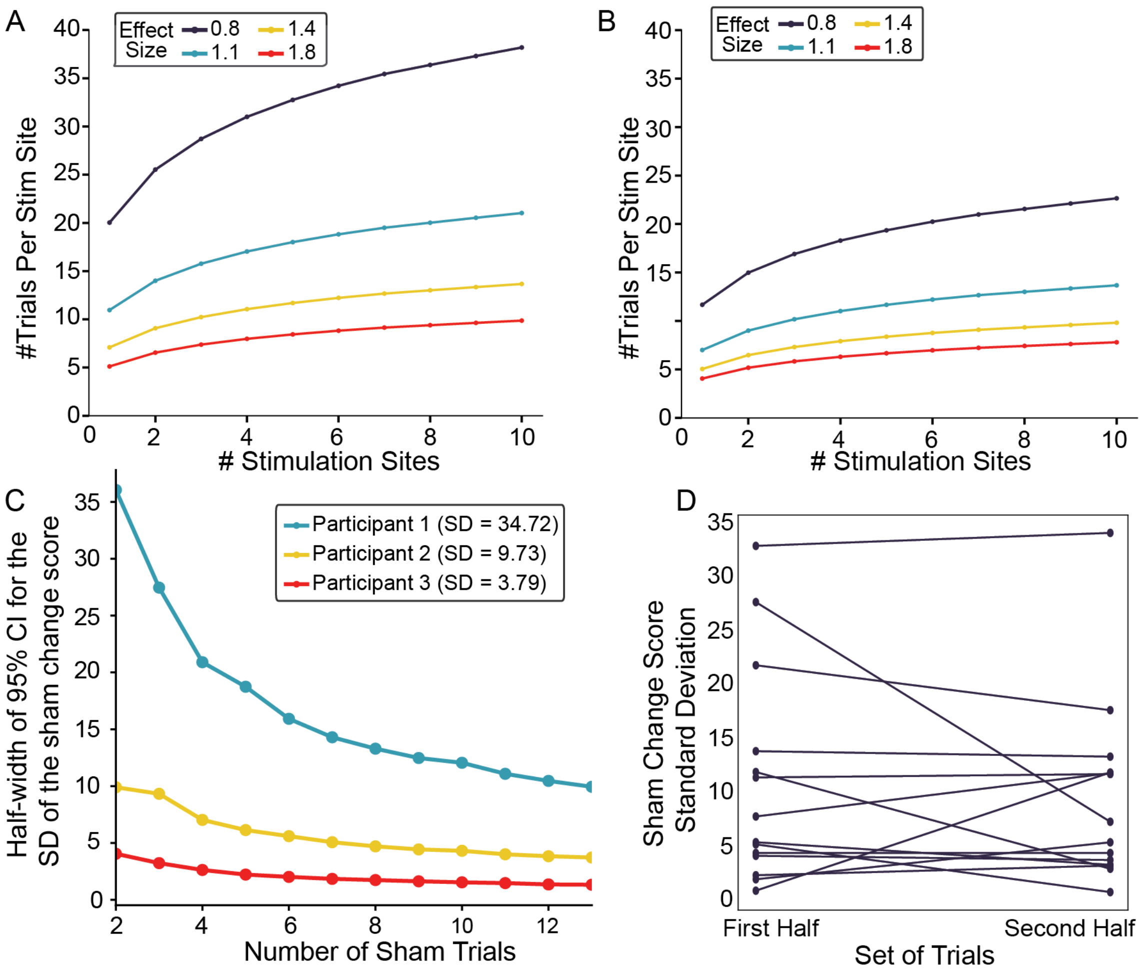


**Supplemental Figure 1. Number of trials required for desired effect sizes in cases of unpaired samples or non-normal distribution**, **determination of the number of sham-only trials required for a stable estimate of sham change score standard deviation, and stability of sham trials across sEEG testing.**

**A:** To find the required sample sizes for unpaired data, statistical power was computed for a one-tailed unpaired t-test with N1/N2 = 1, power = 0.8.

**B:** To find the required sample sizes for a distribution that is not normally distributed, the statistical power was computed for paired signed-rank with power = 0.8.

**C:** Bootstrap stability analysis of sham response variability for three representative participants. The precision of the change score standard deviation estimate, represented by the half-width of the 95% confidence interval (CI), stabilizes after approximately 10 trials for small and moderate standard deviation values.

**D:** Comparison of sham trial change score standard deviation across trials for all participants. The standard deviation of change score sham responses was computed for the first and second half of sham trials. No statistical difference was found between the two sample sets (paired t-test $p = 0.463$, $t = 0.756$; first half [mean (standard deviation)]: 10.51 (9.54), second half: 9.09 (8.26)).

| Stimulation blocks | Sham | Site 1 | Site 2 | Site 3 | Site 4 |
|---|---|---|---|---|---|
| 1 | X | X | | | |
| 2 | X | | X | | |
| . | X | | | X | |
| . | X | | | | X |
| n | X | X | | | |
| A | X | X | | X | X |
| B | X | X | X | X | X |
| C | X | X | X | X | X |
| D | X | X | | X | X |
| E | X | X | X | X | |
| F | X | X | | X | X |
| G | X | X | X | | X |
| H | X | X | X | | |
| I | X | X | X | | |
| J | X | X | X | | |
| K | X | X | X | | X |
| Total Paired Trials | | 13 | 9 | 7 | 8 |

**Supplemental Table 1. Example block design for collecting paired sham and verum stimulation trials.** Preliminary efficacy trials (labeled by number) are collected for a random subset of sites that pass the safety and tolerability testing requirements, as described in Fig 1. Sham trials from preliminary efficacy trials are used to compute expected effect size and estimate the required number of trials per site. Sham-controlled paired testing blocks (labeled by letter) are conducted with sham and verum stimulation. The order of sham and target trials in each block should be counterbalanced (i.e. shuffle the order of Sham and Sites 1-4). For estimates of the required n see supplemental Figure 1C.
